# Development of an exosomal gene signature to detect residual disease in dogs with osteosarcoma using a novel xenograft platform and machine learning

**DOI:** 10.1101/2021.02.11.429432

**Authors:** Kelly M. Makielski, Alicia J. Donnelly, Ali Khammanivong, Milcah C. Scott, Andrea R. Ortiz, Dana C. Galvan, Hirotaka Tomiyasu, Clarissa Amaya, Kristi Ward, Alexa Montoya, John R. Garbe, Lauren J. Mills, Gary R. Cutter, Joelle M. Fenger, William C. Kisseberth, Timothy D. O’Brien, Brenda J. Weigel, Logan G. Spector, Brad A. Bryan, Subbaya Subramanian, Jaime F. Modiano

## Abstract

Osteosarcoma has a guarded prognosis. A major hurdle in developing more effective osteosarcoma therapies is the lack of disease-specific biomarkers to predict risk, prognosis, or therapeutic response. Exosomes are secreted extracellular microvesicles emerging as powerful diagnostic tools. However, their clinical application is precluded by challenges in identifying disease-associated cargo from the vastly larger background of normal exosome cargo. We developed a method using canine osteosarcoma in mouse xenografts to distinguish tumor-derived from host-response exosomal mRNAs. The model allows for the identification of canine osteosarcoma-specific gene signatures by RNA sequencing and a species-differentiating bioinformatics pipeline. An osteosarcoma-associated signature consisting of five gene transcripts (*SKA2, NEU1, PAF1, PSMG2, and NOB1*) was validated in dogs with spontaneous osteosarcoma by qRT-PCR, while a machine learning model assigned dogs into healthy or disease groups. Serum/plasma exosomes were isolated from 53 dogs in distinct clinical groups (“healthy”, “osteosarcoma”, “other bone tumor”, or “non-neoplastic disease”). Pre-treatment samples from osteosarcoma cases were used as the training set and a validation set from post-treatment samples was used for testing, classifying as “osteosarcoma–detected” or “osteosarcoma–NOT detected”. Dogs in a validation set whose post-treatment samples were classified as “osteosarcoma–NOT detected” had longer remissions, up to 15 months after treatment. In conclusion, we identified a gene signature predictive of molecular remissions with potential applications in the early detection and minimal residual disease settings. These results provide proof-of-concept for our discovery platform and its utilization in future studies to inform cancer risk, diagnosis, prognosis, and therapeutic response.

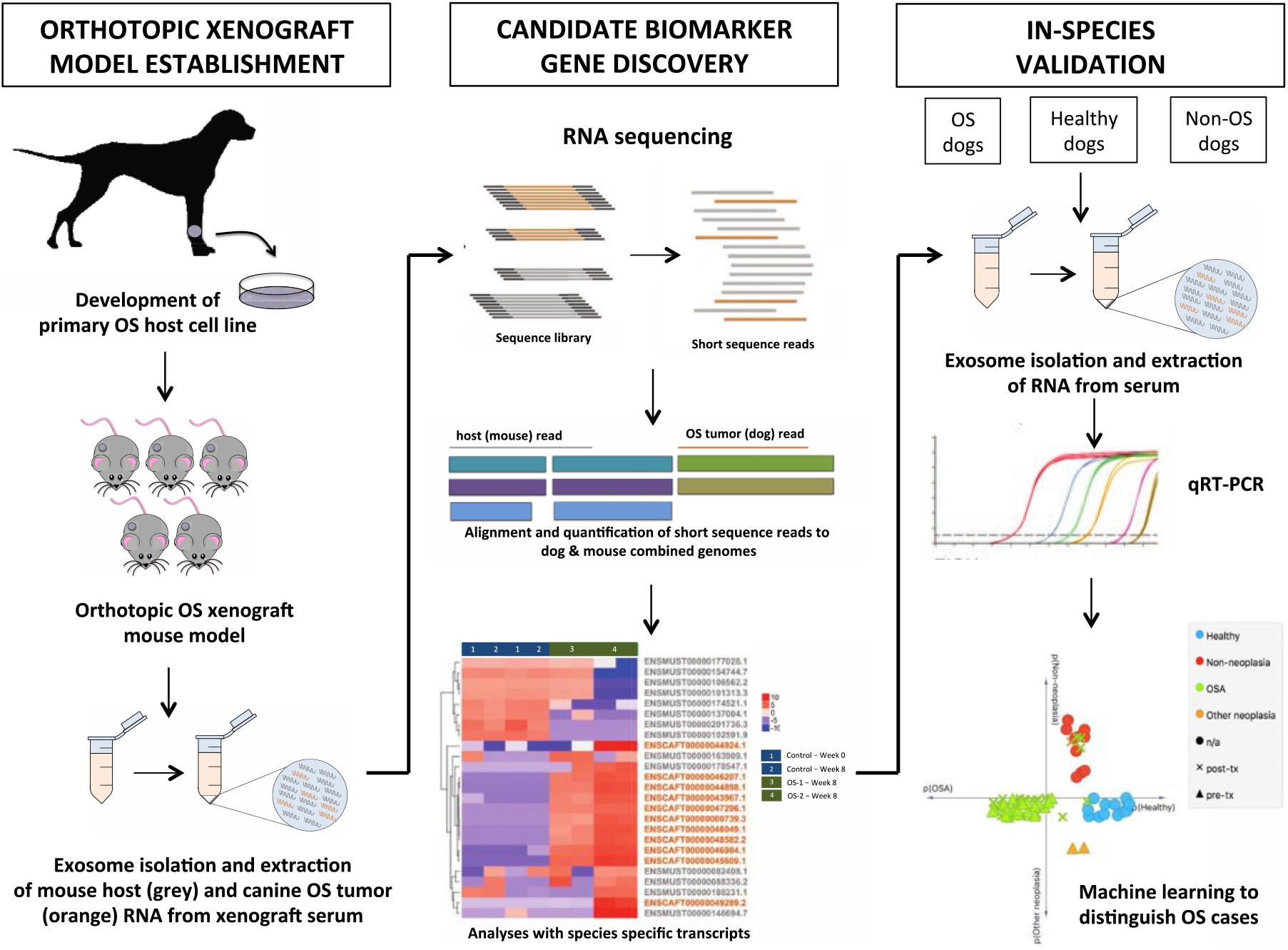

## Introduction

Osteosarcoma is a rare disease that disproportionately affects children, adolescents, and young adults ^1^. More than half of osteosarcoma patients relapse and die from metastatic disease within 10 years of their initial diagnosis ^1,2^, highlighting the need for predictive biomarkers to personalize therapies. Previously, we have identified evolutionarily conserved transcriptional programs with high prognostic value; however, practical obstacles have prevented their wide adoption into clinical practice ^3–5^. Thus, it is apparent that non-invasive tests that inform prognosis and longitudinal remission status represent a continued unmet need for osteosarcoma patients.

Serum exosomes can be used to address these unmet needs in osteosarcoma ^6,7^. Exosomes are secreted, membrane-bound vesicles measuring 30 to 200 nm in diameter that originate from the fusion of multivesicular endosomes to the plasma membrane ^8^. Like other microvesicles, exosomal cargo includes RNA, DNA, proteins, lipids, and cellular metabolites. Exosomes can be powerful diagnostic tools. Specifically, they are stable in biological fluids, can be efficiently and non-invasively isolated, and contain cargo that can be significantly associated with disease states ^9–11^.

Furthermore, the utility of serum exosomes as a diagnostic/prognostic platform is independent of the source and function of such cargo. There have been recent gains in enrichment of exosomes and/or comparably sized microvesicles from blood, plasma, and serum using instrumentation and methodology that is routinely available in diagnostic laboratories ^9–11^, allowing us to envision applications of exosome diagnostics as a realistic goal. However, the identification and differentiation of cargo originating from diseased cells (signal) from the background of normal exosomes (noise) is still a major obstacle that precludes wide use of exosomes in clinical laboratory medicine.

A significant challenge in osteosarcoma research is the development of animal models that accurately represent the complex biology involved in osteosarcoma growth and metastasis. The total number of new human cases diagnosed each year in the United States is low, resulting in a small number of human tumors available for study ^1,5,12^. Osteosarcoma incidence in dogs is approximately 30 times higher than in people; with clinical and molecular evidence to suggest that human and canine osteosarcoma share many key features of tumor organization and disease progression, dogs represent effective models to study these important aspects about the natural history of this disease ^1,13–16^. Gene expression analysis demonstrated conserved, tumor-specific molecular phenotypes in dog and human appendicular osteosarcoma ^5^. This is supported by a more recent study that assessed the transcriptional variation in osteosarcoma tumor samples and cell lines from humans and dogs, demonstrating a high interspecies correlation ^3^.

Previously, we developed a method to identify species-specific messenger RNA (mRNA) sequences in tumor xenografts (tumor, or donor species, and stroma, or host species) ^17,18^. In the present study, we further extended that method to serum exosomes, allowing for identification of a tumor-specific five-gene signature that accurately discriminates osteosarcoma tumor-bearing dogs from dogs in other disease categories and dogs free of apparent disease. Individually, none of the genes could reliably predict the presence of osteosarcoma or minimal residual disease, but when combined with machine learning the five-gene signature could accurately predict the presence of residual disease in dogs undergoing treatment for osteosarcoma. Overall, we demonstrate the discovery of exosome-based biomarkers that have the potential to identify the presence of cancer cells. We also show that this discovery platform that can be used to identify biomarkers to inform prognosis and guide treatment of osteosarcoma in dogs, providing proof of concept to develop and apply comparable approaches for human osteosarcoma patients.

## Materials/Subjects and Methods

### Cell culture

Two canine osteosarcoma cell lines, representing molecular phenotypes associated with different biological behaviors (OS-1 and OS-2), were used in this study^17^. OS-1 and OS-2 are derivatives of the OSCA-32 (Kerafast, Inc., Boston, MA, catalog #EMN003) and OSCA-40 cell lines (Kerafast, Inc., catalog #EMN002), respectively. OS-2 was previously shown to show features associated with aggressive behavior, and OS-1 was shown to behave less aggressively ^17^. Specifically, in an osteosarcoma xenograft model, OS-2 was shown to exhibit a more rapid growth at the primary tumor site, and a greater propensity to metastasize to the lung ^17^. OS-1 and OS-2 cells were modified to stably express green fluorescent protein (GFP; ThermoFisher Scientific, Waltham, MA) and firefly luciferase (luc; ThermoFisher Scientific, Waltham, MA) using a *Sleeping Beauty* transposon system. Cells (1 x 10^6^) were transfected with 1µg of transposase-expressing pDNA vector, *Sleeping Beauty* 100x, and 2 µg of the GFP/luc vector pKT2/CLP-Luc-ZOG in 100 µL of nucleofector solution V (Lonza, Basel, Switzerland, catalog #VCA-1003). Transfected cells were immediately placed into pre-warmed growth medium, and the cells were expanded using Zeocin (Invivogen, San Diego, CA, catalog #ant-zn-05) selection medium. The transfected cells behaved as a unimodal population by flow cytometry. Proliferation and doubling time of the genetically modified cells were comparable to those of the parental cells *in vitro*. GFP-luc transfected cells were used for orthotopic intratibial injections in mice. Prior to mouse injections, cells were grown in exosome-depleted DMEM media containing 5% glucose and L-glutamine (GIBCO, ThermoFisher Scientific, Waltham, MA, catalog #11965) supplemented with 10% exosome-depleted FBS Media Supplement - USA Certified (SBI, Palo Alto, CA, catalog # EXO-FBS-250A-1), 10mM 4-(2-hydroxyethyl)-1-piperazine ethanesulphonic acid buffer (HEPES; ThermoFisher Scientific, Waltham, MA, catalog #15630) and 0.1% Primocin (Invivogen, San Diego, CA, catalog #ant-pm-1), and cultured at 37°C in a humidified atmosphere of 5% CO_2_. Each cell line had been passaged more than 15 times since it was established; however, cell lines were repeatedly authenticated at regular intervals based on short tandem repeats (IDEXX BioResearch, Columbia, MO) to confirm stability during experimentation.

Primary cultures of human pulmonary microvascular endothelial cells (Lonza, Walkersville, MD, catalog #CC-2527) were grown in EGM-2 Endothelial Cell Growth Medium-2 Bullet kit (Lonza). Primary cultures of human pulmonary fibroblasts (Lonza, catalog #CC-2512) were cultured in FGM-2 Fibroblast Cell Growth Medium-2 Bullet kit (Lonza). For exosome depleted conditions, the FBS aliquot in each Bullet kit was excluded from the growth media.

### Exosome purification

Cells were cultured in exosome depleted media and exosomes were isolated using the ExoQuick TC kit (SBI, Palo Alto, CA, catalog #EXOTC10A-1) according to the manufacturer’s instructions.

### Immunohistochemistry and immunofluorescence

Human osteosarcoma tissue microarrays were obtained from US Biomax (Rockville, MD, catalog #OS804), and were staged according to the MSTS staging system. Antigens were detected using immunohistochemistry and quantified by previously described methodology ^19^. For immunofluorescence, cells were treated as indicated and fixed in ice-cold paraformaldehyde solution. Immunofluorescent detection of CD9, CD63, and CD81 was performed using the EXOAB-KIT-1 (SBI, Palo Alto, CA, catalog #EXOAB-KIT-1) and anti-phalloidin-conjugated secondary antibodies. DAPI counter staining was used as a nuclear stain. The anti-human tetraspanin antibodies used for immunohistochemistry and immunofluorescence cross-react against the canine proteins, where each antibody recognizes unique proteins with the correct electrophoretic mobility as determined by immunoblotting.

### Electron microscopy

Exosomes were isolated from OSCA-40 cell culture supernatants using the ExoQuick TC protocol, fixed in 2.5% paraformaldehyde, and washed in PBS. The exosomes were suspended in Milli-Q water, immediately applied to a glass slide, and allowed to air dry for 1 hour. The slides were dehydrated with ethanol, sputtered coated with a gold layer, and imaged using a Zeiss EVO scanning electron microscope.

### Immunoblotting

Immunoblotting was performed to confirm enrichment of tetraspanins (CD9, CD63, and CD81) and concurrent depletion of β-actin, as indicators of the biochemical characteristics of exosomes reported in the literature. 200µl of Pierce RIPA buffer (ThermoFisher Scientific, Waltham, MA, catalog #89900) combined with 1µl of HALT protease inhibitor and 1µl of HALT phosphatase inhibitor cocktail (ThermoFisher Scientific, Waltham, MA, catalog #78420) were added to cell or exosome pellets and vortexed for ∼15 seconds. The samples were incubated at room temperature for 5 minutes to allow complete lysis before pre-clearing nuclei and insoluble material by centrifugation. The protein content was determined using the BCA assay kit as recommended by the manufacturer (ThermoFisher Scientific, Waltham, MA, catalog #23225). For immunoblotting, 50µg of protein for each sample were diluted into Laemmli buffer, heated at 95⁰C, and then immediately chilled on ice before loading onto the gels. SDS-PAGE electrophoresis and transfer to PVDF membranes was done using routine protocols ^20^. Membranes were blocked with 5% dry milk in Tris Buffered Saline containing 0.05% Tween (TBS-T), followed by overnight incubation at 4°C with antibodies directed against CD9, CD63, and CD81 (SBI, Palo Alto, CA, catalog #EXOAB-KIT-1) at 1:1000 dilution in TBS-T buffer containing 5% dry milk. Anti-β actin was used as described ^21^ to serve as a control for depletion of cytosolic proteins in exosomes. Blots were washed and incubated for one hour at room temperature with a secondary goat anti-rabbit-HRP antibody at 1:20,000 dilution. The blots were finally incubated with chemiluminescence substrate and visualized on a LI-COR Odyssey Imager (LI-COR, Lincoln, NE).

### Nanoparticle tracking

Exosomes were enriched from samples as detailed above and resuspended in PBS to a total volume of 1mL. The size distribution of extracellular vesicles was measured using a NanoSight Nanoparticle Tracking Analyzer (Salisbury, United Kingdom), using the settings recommended by the manufacturer. Size, frequency, and distribution measurements were recorded in triplicate for each sample and were analyzed by the built-in NanoSight Software.

### Plasmids and transfection

Transduction of osteosarcoma cells with genes encoding tetraspanins, which are enriched in exosomes, was performed using the pCT-CMV-GFP-MCS-EF1α-Puro lentiviral system (SBI, Palo Alto, CA, catalog #CYTO800-PA-1) as described ^22^. Puromycin was used to select for stably transfected cells, and cells were grown in exosome depleted media prior to exosome collection.

### Tumor xenografts

Six week-old, female, athymic nude mice (strain NCr nu/nu) were obtained from Charles River Laboratories (Wilmington, MA). Animals were assigned to separate cages in random order for each experiment. All mouse experiments were approved by The University of Minnesota Institutional Animal Care and Use Committee (Protocols 1307-30806A, 1606-33857A, and 1803-35710A). For intratibial (I.T.) injections, mice were anesthetized with xylazine (10 mg/kg, intraperitoneally (I.P.)) and ketamine (100mg/kg, I.P.) in preparation for orthotopic I.T. injections. Canine osteosarcoma cells were suspended in sterile PBS (ThermoFisher Scientific, Waltham, MA, catalog #10010049) and 10 µL containing 1 x 10^5^ cells were injected I.T. as previously reported ^17,21^. Control mice had 10 µL sterile PBS injected I.T. All injections were administered into the left tibia using a tuberculin syringe with 29-gauge needle. For each osteosarcoma cell line, OS-1 and OS-2, five mice received cell-I.T. injections; 3 mice received sham (PBS)-I.T. injections. Buprenorphine (0.075mg/kg, I.P. every 8 hours) (Reckitt Benckiser Healthcare, Richmond, VA) was administered for analgesia for 24 hours following the injections, and prophylactic ibuprofen was administrated in the water for the next 3 days. Mice were monitored by weekly bioluminescence imaging and tumor size measurements. At 8 weeks after the injections, the mice were humanely euthanized using a barbiturate overdose. Blood was collected via intracardiac phlebotomy. The tibiae and the lungs were collected from mice injected with osteosarcoma cells (n = 5 for each cell line) and placed in 10% neutral buffered formalin for histopathology or stored at −80°C. The presence of tumors was confirmed histologically.

### Osteosarcoma xenograft serum exosome enrichment and RNA extraction

Exosomes were enriched from serum samples from control mice and from tumor bearing mice at week-8 using ExoQuick reagent (SBI, Palo Alto, CA, catalog #EXOQ5A-1) according to the manufacturer’s instructions. Briefly, serum was mixed with ExoQuick reagent at a volume of 252 µL ExoQuick per 1 mL of serum. The mixture was incubated for 30 minutes at 4°C, followed by centrifugation at 1,500 x g for 30 minutes to enrich exosomes. The resulting supernatant was discarded, and the tubes were centrifuged for an additional 5 minutes at 1,500 x g to remove any remaining supernatant. Exosomal RNA was extracted using SeraMir ExoRNA Amp Kit (SBI, Palo Alto, CA, catalog #RA800A-1), according to the manufacturer’s instructions.

### Library preparation and next-generation sequencing

Pooled serum from each group was sequenced and analyzed. Sequencing libraries were prepared using the Clontech SMARTer® Stranded Total RNA-Seq Kit v2 - Pico Input Mammalian kit (Takara Bio, Kasatsu, Japan). RNA sequencing (50-bp paired-end, with HiSeq 2500 Illumina) was performed at the University of Minnesota Genomics Center (UMGC). A minimum of 16 million read-pairs was generated for each sample and the average quality scores were above Q30 for all pass-filter reads.

### Bioinformatics analysis

Initial quality control analysis of RNA sequencing FASTQ data was performed using FastQC software (v0.11.5). FASTQ data were trimmed with Trimmomatic (v0.33.0). Kallisto (v0.43.0) was used for pseudoalignment and quantifying transcript abundance. For accurate alignment of sequencing reads to canine and murine genes, a kallisto index was built from a multi-sequence FASTA file containing both the canine (CanFam3.1) and murine (GRCm38.p5) genomes. For each species, transcripts <200bp were removed from the FASTA files. The masked FASTA files were then merged for a total of 121,749 murine and canine transcripts. Insertion size metrics were calculated for each sample using Picard software (v1.126). Data will be deposited in GenBank/GEO. The ‘DESeq2’ package in RStudio was used for differential analysis of transcript counts obtained from kallisto. Transcript counts were first summarized to gene counts and then DESeq2 was used to convert count values to integer mode, correct for library size, and estimate dispersions and log2 fold changes between comparison groups. Genes with a Benjamini-Hochberg adjusted p-value < 0.05 and log2 fold change >+/-4 between control and xenograft samples were considered significantly differentially expressed transcripts. Statistically differentially expressed canine genes were removed if they had a DESeq2 normalized value of greater than zero in the control group (mouse sequences) as these would be highly homologous genes between the mouse and dog. Counts per million (CPM) values of genes were log2 transformed and mean-centered prior to clustering. The ComplexHeatmap package was used for clustering and creating heatmap figures. Enriched pathway and functional classification analyses of differentially expressed transcripts were performed using QIAGEN’s Ingenuity® Pathway Analysis (IPA®; QIAGEN, Redwood City, CA). The reference set for all IPA analyses was the Ingenuity Knowledge Base (genes only). Canine-associated gene names were used as the output format for input datasets with canine genes and murine-associated gene names were used as the output format for input datasets with murine genes.

### qRT-PCR validation of sequencing data

Serum or plasma samples were obtained from client-owned dogs with naturally-occurring osteosarcoma before and after treatment as part of routine biobanking efforts. The samples included in the analysis were identified retrospectively. Serum samples were also obtained and biobanked from client-owned dogs that were hospitalized with various non-malignant conditions. For healthy controls, serum samples were obtained from staff- and student-owned dogs with no apparent disease. The samples were divided into a training set and a test set. The training set included dogs in one of four categories (“osteosarcoma”, “other neoplasia”, “non-neoplasia”, and “healthy” (dogs with no apparent disease)). Osteosarcoma dogs had a primary tumor of bone and were treatment naïve (**Table 1**). The test set included samples from dogs with osteosarcoma, after treatment (n = 24; **Table S1**). Blood was collected into vacutainer tubes that were centrifuged at 3,000 x g for 15 minutes. Aliquots of serum or plasma were transferred to 1.5 ml microcentrifuge tubes and stored at −80°C until analysis. All treatment decisions were at the discretion of the attending clinician. All procedures were approved by the Institutional Animal Care and Use Committees of The University of Minnesota under protocols 0802A27363, 1101A94713, 1312-31131A, 1504-32486A, 1702-34548A, 1803-35759A, and 2003-37952A and The Ohio State University 2010A0015-R2 and 2018A00000100. Exosomes were precipitated from canine serum or plasma samples using ExoQuick serum reagent (SBI, Palo Alto, CA, catalog #EXOQ5A-1) according to the manufacturer’s instructions. Additional steps were included for plasma samples: 10 µL of thrombin (SBI, Palo Alto, CA, catalog #TMEXO-1) was added for each 1 mL of plasma. The sample was then mixed at room temperature for 5 minutes, followed by centrifugation at 10,000 rpm for 5 minutes. The supernatant was transferred to a new microcentrifuge tube, and the volume recovered was noted. Plasma and serum samples were subsequently treated the same. Briefly, the sample was mixed with ExoQuick reagent at a volume of 252µl ExoQuick per 1ml of serum. The mixture was incubated for 30 minutes at 4°C, followed by centrifugation at 1,500 x g for 30 minutes to precipitate exosomes. The resulting supernatant was removed and discarded, and the tubes were centrifuged for an additional 5 minutes at 1,500 x g to remove any remaining supernatant. Exosomal RNA was extracted using the mirVana miRNA Isolation Kit (Ambion, Thermo Fisher Scientific, Waltham, MA), according to the manufacturer’s instructions. Elimination of genomic DNA and reverse transcription were both carried out using QuantiTect Reverse Transcription Kit (Qiagen, Valencia, CA). Real-time quantitative reverse transcription PCR (qRT-PCR) was performed on a LIGHTCYCLER 96 (Roche, Indianapolis, IN) with FastStart SYBR Universal Green Master Mix (Roche, Indianapolis, IN) Protocol. GAPDH was used as the reference standard for normalization ^23^ and relative levels of steady state mRNA were established using the comparative [delta]Ct method. The relationship between RNA-sequencing data and qRT-PCR values for the transcripts of interest were analyzed using Pearson’s correlation.

**Table 1.**
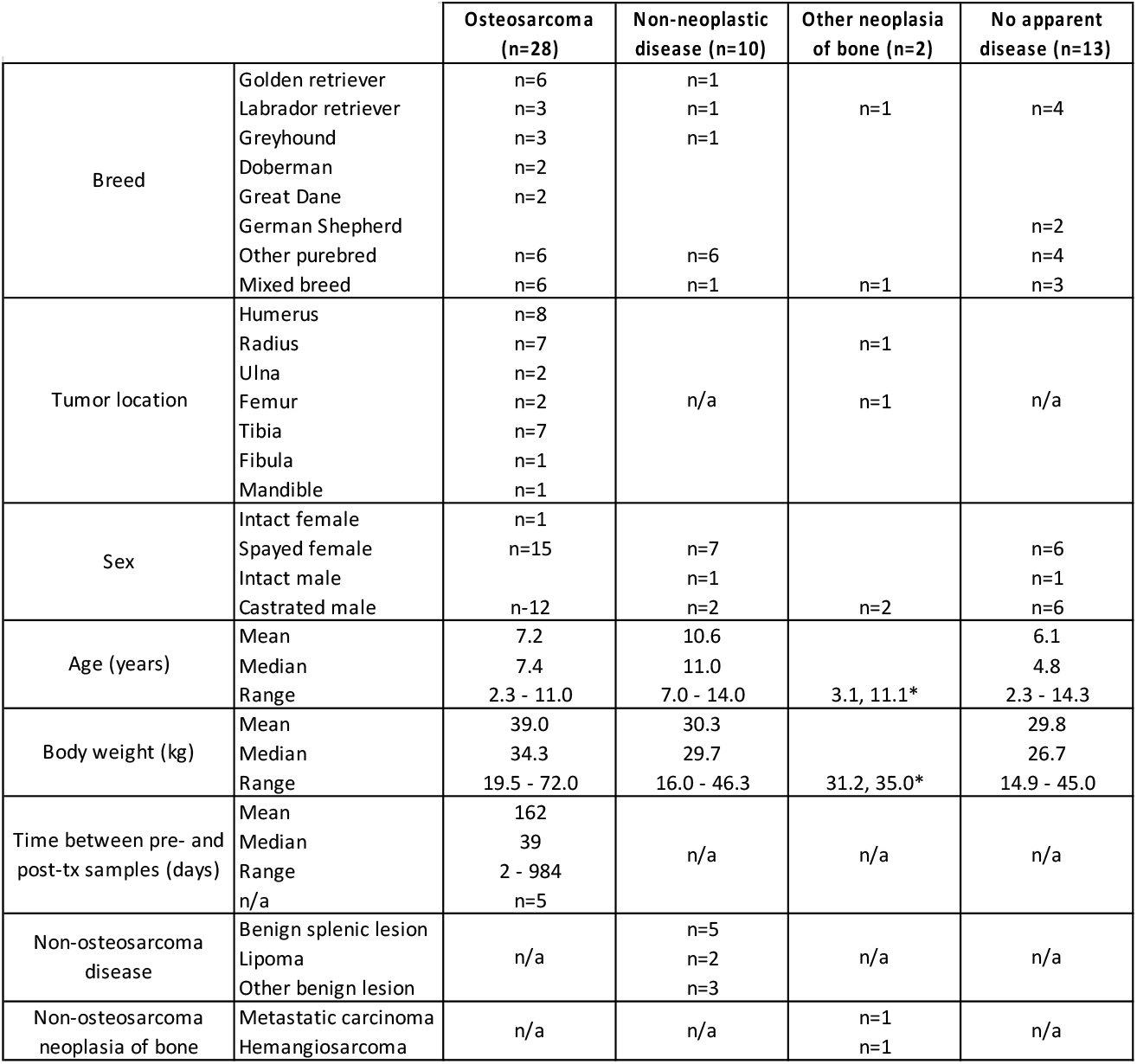
Clinical characteristics of enrolled dogs. * Mean and median values are not listed for the “Other neoplasia of bone” category, as only two dogs were enrolled in this group.

### Machine learning

Gene expression data from samples of dogs with “no apparent disease” (healthy, n=13), non-neoplastic/benign conditions treated with surgery (diseases other than cancer; n=10), osteosarcoma (n=27), and other neoplasia (non-osteosarcoma cancers; n=2) pre-treatment samples (52 total) normalized to GAPDH (as internal control) were used to train and build different machine learning models. The normalized data were further transformed using three-component linear discriminant analysis (LDA). Different machine learning algorithms were then tested and compared to identify the top-performing predictive models that fit well with our data, including Logistic Regression (LR), Linear Discriminant Analysis (LDA), k-Nearest Neighbors (KNN), Decision Tree (CART), Gaussian Naïve Bayes (NB), Support Vector Machine (SVM), Bagging (BAG), Random Forest (RF), Extra Trees (EXT), Adaptive Boosting (ADA), Stochastic Gradient Boosting (SGB), Neural Network (NN), Ridge regression (RGD), and Stochastic Gradient Descent (SGD) classifiers Scikit-learn Python package (http://scikit-learn.sourceforge.net) ^23^. For training and optimization, the training dataset was randomly split into training and validation sets using k-fold cross-validations with sample stratification (when possible). k-fold cross-validation randomly splits data into k groups, where k - 1 groups were used for training and one remaining group was used for validation; repeated for k times with each of k validation sets being used only once. The k-fold cross-validation was then repeated and averaged across 100 iterations with random shuffling in between to ensure performance stability across multiple tests. For this study, a 10-fold cross-validation was used. Top models with the best averaged sensitivity and specificity were chosen for further optimization and testing. The sensitivity was calculated based on the equation True Positives / (True Positives + False Negatives) and the specificity was True Negatives / (True Negatives + False Positives)^24^. True positives were defined as the classification accuracy for osteosarcoma and true negatives were defined as the classified accuracy of non-osteosarcoma. Predictive power was also estimated for the final top-performing models based on their positive (PPV) and negative (NPV) predictive values. The PPV was calculated as True Positives / (True Positives + False Positives) and NPV was True Negatives / (True Negatives + False Negatives) ^24^. Data from the unknown samples (post-treatment osteosarcoma subjects) were transformed based on the fitted training set and classified using the top trained learning models. The resulting classification calls were further tested against survival data of the post-treatment osteosarcoma subjects over time as a means for establishing the significance that (detectable) minimal residual disease had on event-free survival times.

Post-treatment osteosarcoma samples classified as “osteosarcoma” by all of the selected top machine learning models (either most accurate or most sensitive) were considered to be “osteosarcoma-detectable”. Post-treatment osteosarcoma samples that received another classification by one or more of the top models were considered to be “osteosarcoma-NOT detectable”. Kaplan-Meier survival analysis was performed using R packages *survival* (v3.27) and *survminer* (v0.48). A log-rank (Mantel-Cox) test was used to compare event-free survival times between dogs whose post-treatment samples were considered “osteosarcoma-detectable” and dogs whose post-treatment samples were considered “osteosarcoma-NOT detectable”.

## Results

### Exosome production by osteosarcoma cells is positively correlated with tumor aggressiveness

Previous studies have documented a quantitative relationship between tumor aggressiveness and the amount of exosomes produced ^11^. To investigate this in the context of osteosarcoma, the presence of tetraspanins CD9 and CD63, two transmembrane proteins that are enriched in exosomes and other microvesicles, was quantified in 80 human osteosarcoma samples using immunohistochemistry (IHC) (**Figs. 1A-B**). The data show that stage III tumors stained more robustly for both CD9 and CD63 than stage I or stage II tumors (**Figs. 1A-B**), suggesting that a positive relationship between total detectable exosomes or exosomal protein and tumor stage also exists in osteosarcoma. To further delineate the functional involvement of exosomes in the progression of osteosarcoma, we sought to use a more tractable *in vivo* model system for exosome biomarker discovery. Given the molecular and biological similarities between human and canine osteosarcomas ^12,16^, a spontaneous canine model was used for our studies. Our immediate next experiments were thus devoted to characterizing canine osteosarcoma-derived exosomes and to confirm their conserved roles in the biology of the disease ^6,7^. We first validated exosome production by canine osteosarcoma cell lines using immunofluorescence (IF). Previous experiments showed that the biologic behavior of the primary tumors was conserved in the cell lines, with OSCA-40 being more aggressive, as it formed tumors that grew rapidly and showed a greater propensity for pulmonary metastasis than two counterparts, called OSCA-32 (a.k.a., OS-1) and OSCA-8, in an orthotopic xenograft model ^17,21^. **Fig. S1** shows positive staining for CD63, CD9 and CD81 in secreted microvesicles from OSCA-32 and OSCA-40 canine osteosarcoma cell lines by IF, and was further confirmed by immunoblotting (**Fig. 2B**, and data not shown). Our results indicate the conservation of tetraspanins in extracellular vesicles, most likely representing exosomes, from humans and dogs.

**Figure 1.**
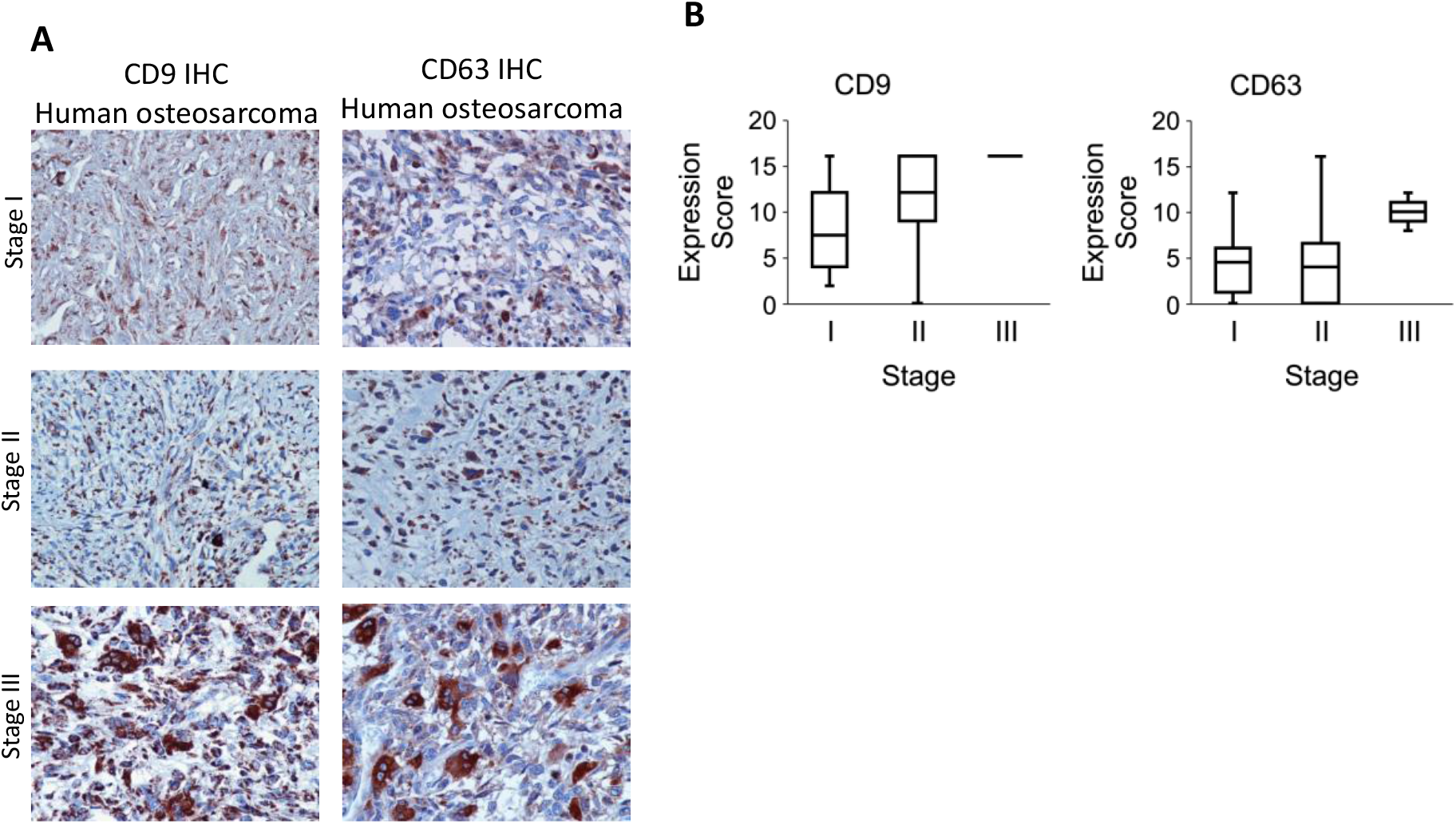
Exosome production by osteosarcoma cells is positively correlated with tumor progression. (A) Representative images from exosome specific staining in human osteosarcoma tissues. Brown staining indicates positive detection of exosomes. (B) Tissue biopsy samples from human osteosarcoma patients were stained for the presence of exosome markers CD9 and CD63. Box and whisker plots indicate the IHC expression level of the two exosome markers across stage I, II, and III osteosarcomas.

**Figure 2.**
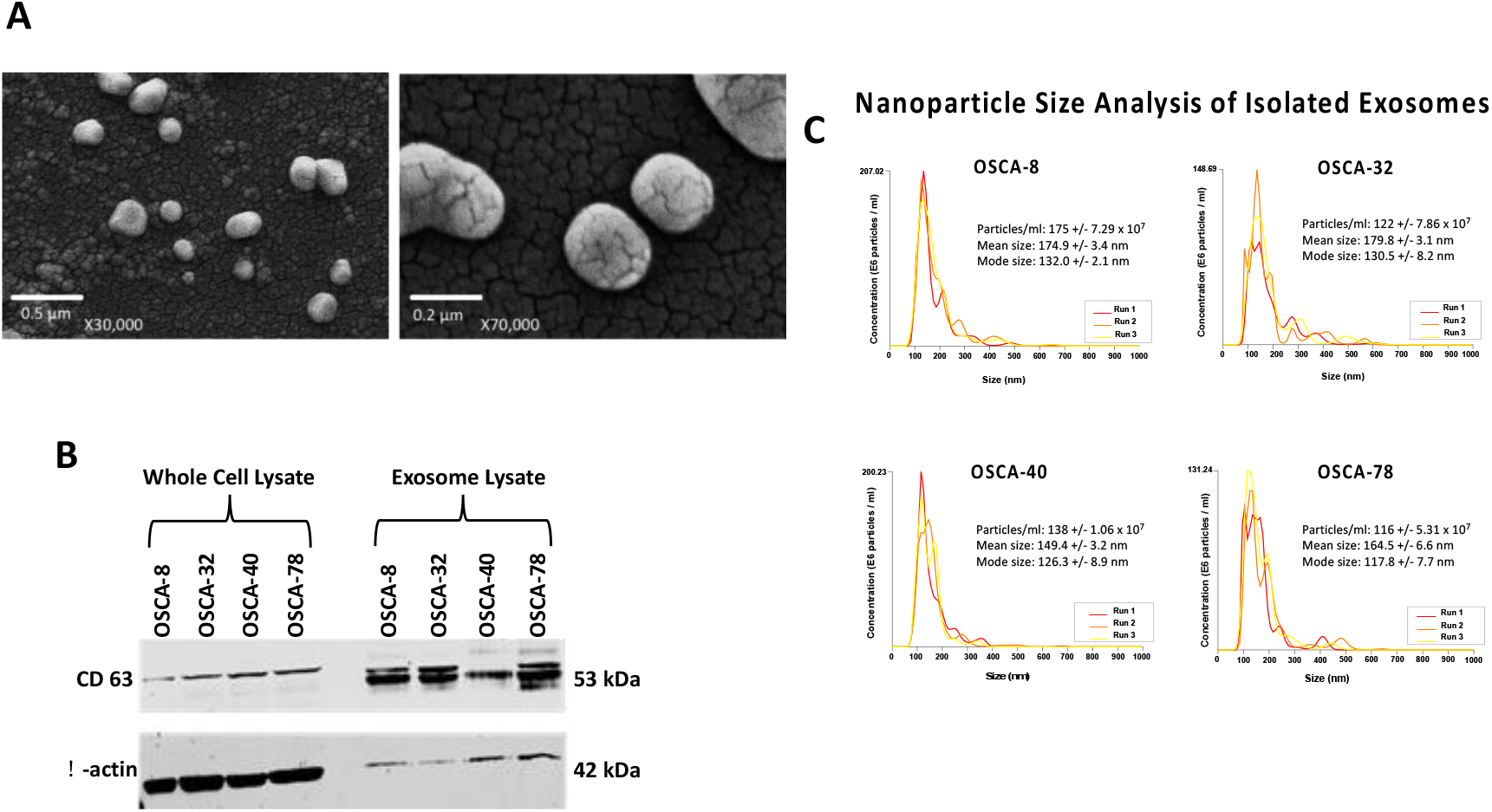
Physical and biochemical characterization of osteosarcoma exosomes. (A) Scanning electron micrographs of OSCA-40 exosomes (B) Immunoblotting of exosome preparations documenting enrichment of tetraspanins (CD63) and depletion of β-actin in OSCA-8, OSCA-32, OSCA-40, and OSCA-78 dog osteosarcoma cell lines. (C) NanoSight particle tracking analysis of triplicate samples each of osteosarcoma cell lines OSCA-8, OSCA-32, OSCA-40, and OSCA-78, showing modal diameters of approximately 130 nm.

We then aimed to confirm the physical properties of canine osteosarcoma-derived extracellular vesicles as exosomes, such as enrichment of tetraspanins (CD9, CD63, and CD81) with concurrent depletion of β-actin, as well as their shape, structure, and size. We utilized scanning electron microscopy (SEM), NanoSight particle tracking analysis, and immunoblotting to validate exosome enrichment from cell lines and from serum (**Figs. 2A-C** and **Fig. S2**). SEM showed spherical microvesicles between 100 and 200 nm in diameter; this size and shape was consistent with that predicted for exosomes (**Fig. 2A**). Additionally, immunoblotting showed enrichment of CD63 and depletion of β-actin in the osteosarcoma cell line-derived exosomes relative to the whole cell lysates (**Fig. 2B**). Exosomal enrichment of tetraspanins, including CD9, CD63, and CD81, with concurrent depletion of β-actin, is consistent with what is reported in the literature for exosome validation ^8^. Finally, nanoparticle tracking analysis showed that the mean vesicle size ranged from 149 nm – 180 nm with a mode of 117 nm – 132 nm, (**Fig. 2C**). This range is similar to the microvesicle size determined by SEM and is also consistent with the expected size of exosomes ^25–32^. We also confirmed that exosomes enriched from serum samples of a dog with osteosarcoma and a dog with no evidence of disease have size distributions comparable to cell line-derived exosomes, as determined by nanoparticle tracking (**Fig. S2**). In all, these findings support the methodology used for exosome enrichment from cell lines and from serum samples.

### Osteosarcoma-derived exosomes can be internalized by stromal cells to modulate gene expression and induce invasive cell behavior

The high propensity for distant metastatic growth in both human osteosarcoma patients and dogs with osteosarcoma has been well documented and is a key factor in survival rates ^1,2,33^. The importance of exosomes in promoting a pre-metastatic niche has been characterized in pancreatic cancer ^34,35^ as well as melanoma ^36^. However, the ability of osteosarcoma exosomes to influence the complex cascade of events that occurs during metastasis, particularly their impact on stromal cells within the microenvironment, is still being determined. Baglio *et al*. documented the ability of osteosarcoma-tumor extracellular vesicle-educated mesenchymal stem cells (MSCs) to promote tumor growth and metastasis in an orthotopic xenograft mouse model of osteosarcoma ^37^. The importance of cancer-associated fibroblasts and endothelial cells in tumor progression has been well described ^38^, making them a useful model system to investigate exosome internalization and influence on cell behavior. Ascertainment that secreted tumor-derived exosomes can be taken up by stromal cells in the organ that is the major target of metastasis requires a method to track tumor specific exosomes. This was accomplished by transfecting canine osteosarcoma cells with CD81 linked to a green fluorescent protein (GFP) tag.

Expression of the fusion protein in transfected OSCA-40 cells, and its incorporation into secreted microvesicles, were visualized using IF microscopy (**Fig. S3A**). Lung stromal fibroblasts and lung endothelial cells were selected as the most relevant target cells for analysis. **Figs. S3B-E** show that GFP-positive osteosarcoma-derived exosomes were internalized by human pulmonary fibroblasts or human pulmonary endothelial cells within 6 to 8 hr, with nearly 100% of the target cells showing GFP expression by 24 hr.

### Serum-derived exosomal canine gene signatures identified in mouse xenografts are associated with osteosarcoma

The *in vitro* data suggested that exosomes influence the tumor microenvironment; however, it remained unclear if these studies were directly translatable to *in vivo* studies where tumor cells maintain a series of complex relationships within their local environment. To address the concerns of *in vitro* translatability, a previously described orthotopic intratibial xenograft mouse model was utilized ^17,21^. Briefly, we established xenografts in nude mice using two canine osteosarcoma cell lines with different metastatic propensities, collected serum exosomes from these mice and from sham-treated controls (injected intratibially with PBS) and performed next-generation sequencing to characterize the full complement of exosomal mRNAs derived from the tumors, as well as from the host response ^17^. Predictably, no xenograft (canine) mRNAs were detectable in sera from the mice prior to tumor implantation, but canine mRNAs were readily apparent in sera from mice with established tumors (**Fig. 3A**). Interestingly, the exosomal transcripts identified in cultured canine osteosarcoma cells showed only minimal overlap (1.4%) with the exosomal transcripts derived from the same canine cell lines when they formed tumors *in vivo* (data not shown), suggesting that the microenvironment is a major factor influencing exosome loading. Further analysis identified groups of canine exosomal mRNAs with correlation scores >0.8 that were part of canonical signaling pathways including cell death, cell signaling, metabolism, and immune response. Changes in the mRNA content of host exosomes were also detectable. Thirty-eight differentially expressed mouse mRNAs were identified in exosomes from animals bearing xenografts when compared to the sham-treated controls (**Fig. 3B**). These mRNAs were primarily associated with immune signaling, including IL-12 signaling, and cellular metabolism (**Fig. 3C**), as well as macrophage activation and stromal cell activity. Intriguingly, atherosclerosis signaling pathways, as well as cholesterol and fat pathways (LXR, FXR) were also identified; however, we suspect this is related to the activation of macrophage-driven processes.

**Figure 3.**
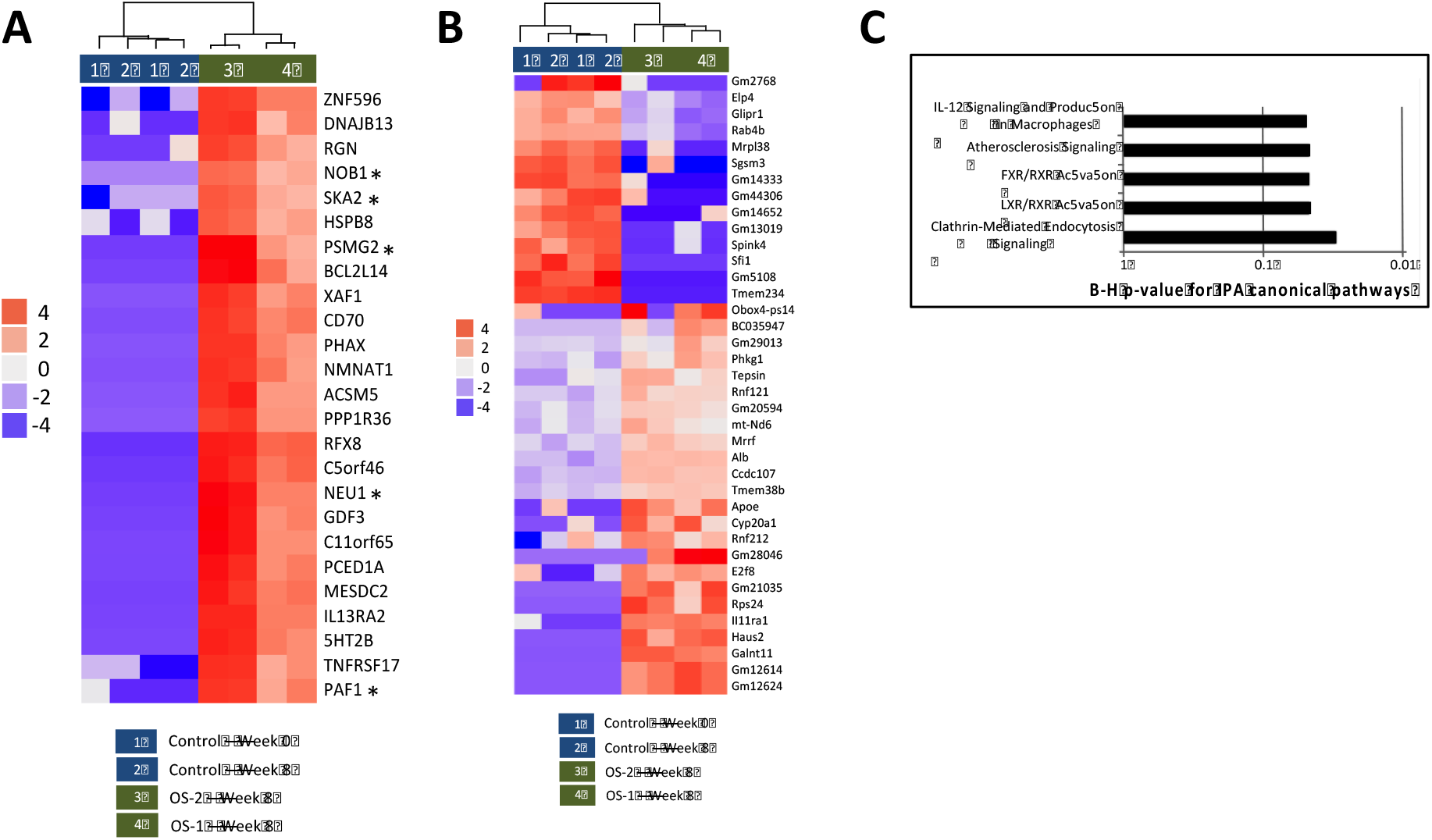
Detecting biomarkers of disease and host response. Heatmap of 25 differentially expressed dog transcripts (A) and 38 differentially expressed mouse transcripts (B) identified by statistical testing with ‘DESeq2’. Colored toe bars represent the different experimental samples. Asterisks in (A) indicate the genes incorporated into the osteosarcoma gene signature. The color coded (red to blue) scale represents +/− change gene expression. (C) Pathways identified by Ingenuity® Pathway Analysis as being associated with differentially expressed host (mouse) genes.

### A five-gene “osteosarcoma-detectable” signature predicts prognosis in dogs with osteosarcoma after treatment

After validating the enrichment of exosomes with tetraspanins, we then looked for differences within exosome cargo that allowed for the determination of an osteosarcoma-specific gene signature. The absence of biomarkers to guide treatment is a major obstacle that has hindered progress in osteosarcoma therapy for dogs and humans alike. <u>We believe that clusters of co-expressed exosomal mRNAs could provide such biomarkers, independent of their biological function.</u> Twenty-five canine mRNAs were reproducibly identifiable and highly expressed in tumor-derived serum exosomes (*i.e.*, were always present in sera from mice with canine osteosarcoma tumors, but not in sera from sham-treated mice, **Fig. 3A**). To build a diagnostic biomarker set, we narrowed the list to the 10 mRNAs with the highest expression, with low inter-sample variation. Five of the 10 mRNAs, representing transcripts from the *SKA2, NEU1, PAF1, PSMG2, and NOB1* genes were determined to be suitable candidates for the biomarker set by virtue of being detectable in serum exosomes derived from dogs with osteosarcoma.

“*In-species* validation” was done by evaluating abundance of these five exosomal mRNAs in 80 archival serum and/or plasma samples from 53 dogs divided into four groups. The clinical characteristics of enrolled dogs are shown in **Table 1**. Serum or plasma samples were included as available in sample archives, and plasma samples were treated with the addition of thrombin to precipitate clotting factors; thereafter the samples were treated similarly, as detailed in the methods. Serum and plasma samples were available for simultaneous testing in a limited number of cases, and the results were concordant. The first group consisted of 28 samples from dogs with osteosarcoma. Of these, 26 included serum or plasma collected prior to and at various time points after treatment (amputation +/− chemotherapy) ranging from 2 to 984 days (median = 37), one only included serum collected before treatment, and one only included serum collected after treatment. The second group consisted of 10 samples from dogs that had been diagnosed with non-neoplastic diseases. The third group consisted of two samples from dogs with intramedullary soft tissue sarcomas (metastatic carcinoma and hemangiosarcoma, the latter of which had pre- and post-treatment samples). The fourth group consisted of 13 samples from dogs with no apparent disease, included as unaffected controls (henceforth referred to as “healthy”) (**Table 1**).

**Figs. 4A-E** show the relative abundance of each of the five mRNAs in the biomarker set, measured by qRT-PCR, in serum exosomes from each subgroup of dogs; **Fig. 4F** shows the relative contribution of each of these mRNAs based on the F-values from the analysis of variance (ANOVA) as applied to the machine learning training algorithms described below.

**Figure 4.**
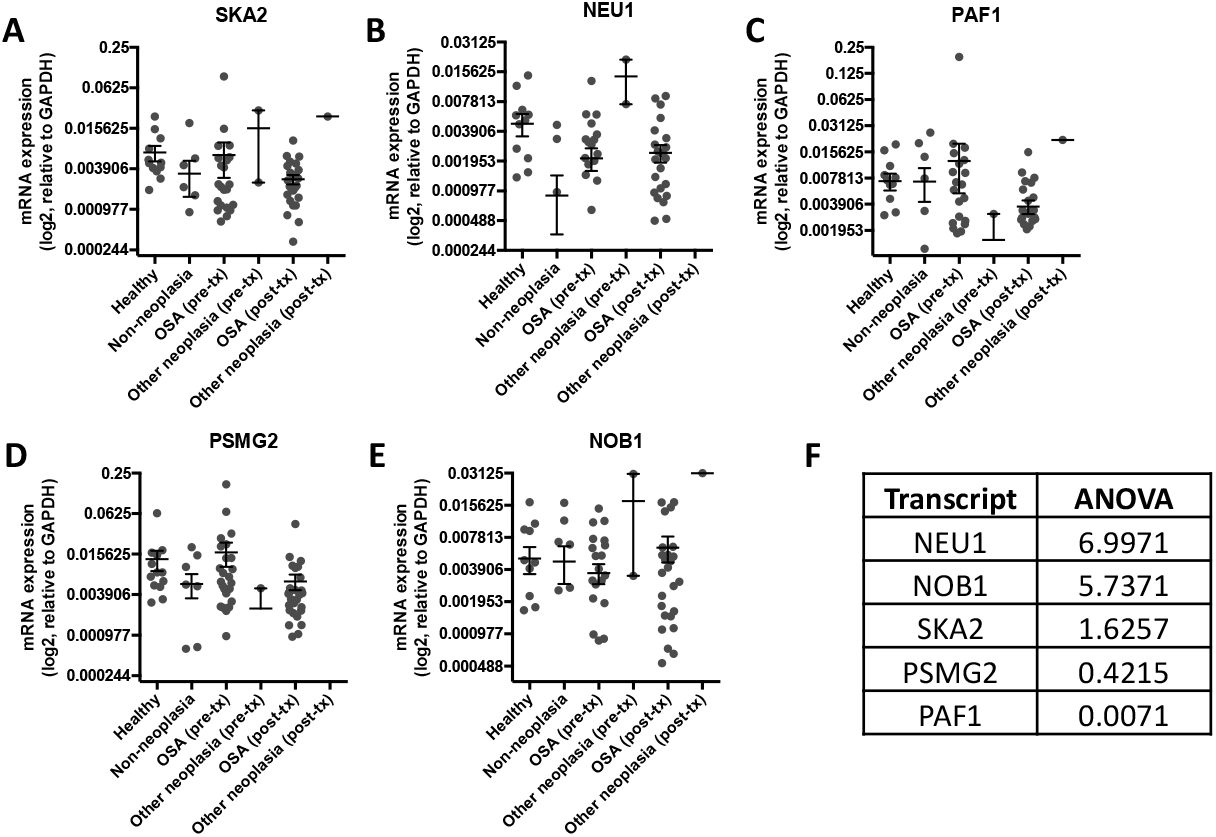
Genes identified in xenografts were validated in canine serum-derived exosomes. Gene expression analysis of (A) SKA2, (B) NEU1, (C) PAF1, (D) PSMG2, and (E) NOB1 gene transcripts by qRT-PCR. Relative expression data were mean centered and scaled to the standard deviation across all samples for each gene. (F) Table showing ANOVA values, demonstrating the relative contribution of each gene transcript to the overall gene signature.

To develop a predictive diagnostic tool to identify dogs with osteosarcoma and distinguish osteosarcoma from the other conditions under test, we built and compared different machine learning models using 53 samples as a training set. These samples included those from “healthy” dogs, dogs with non-neoplastic diseases (“non-neoplasia”), dogs with intramedullary soft tissue sarcomas (“other neoplasia”) and dogs with osteosarcoma prior to treatment (“pre-treatment osteosarcoma”) as a training set to build machine learning models. Stratified 10-fold cross-validation analysis of the training set was performed across 14 different machine learning algorithms based on the five-gene features combined with 3-component LDA-transformation and repeated for 100 iterations with shuffling in between (**Fig. 5**). The top-performing models based on sensitivity and specificity were k-nearest neighbors (KNN), bagging (BAG), random forest (RF), and extra trees (EXT) classifiers (**Fig. 5**, red dashed boxes). The three most sensitive models were logistic regression (LR), linear discriminant analysis (LDA), and ridge (RDG) classifiers (**Fig. 5**, blue dashed boxes). The mean sensitivity (the prediction accuracy for “osteosarcoma”) for the top-performing models ranged from 72 – 82%, while the mean specificity (“non-osteosarcoma”) was between 44% and 51%. The lower specificity was largely due to poor classification of “non-neoplasia” and “other neoplasia” groups as compared to “healthy” and “osteosarcoma” (**Fig. S4**). When the prediction for “healthy” (*i.e.*, dogs with no apparent disease) was analyzed independently, the mean specificity was indeed higher at around 60% for the top four models (**Fig. S4A**), while the prediction accuracy for “non-neoplasia” and “other neoplasia” was only between 0 and 2% **(Fig. S4B).** While the mean specificities for the most sensitive models (LR, LDA, and RDG) were below 25%, their mean sensitivities were ≥ 89%. To show that the results were dependent on the relevant groups in the training set, group assignments were randomized. The machine learning performance was greatly affected following data randomization, showing a significant reduction in both sensitivity and specificity **(Fig. S5).**

**Figure 5.**
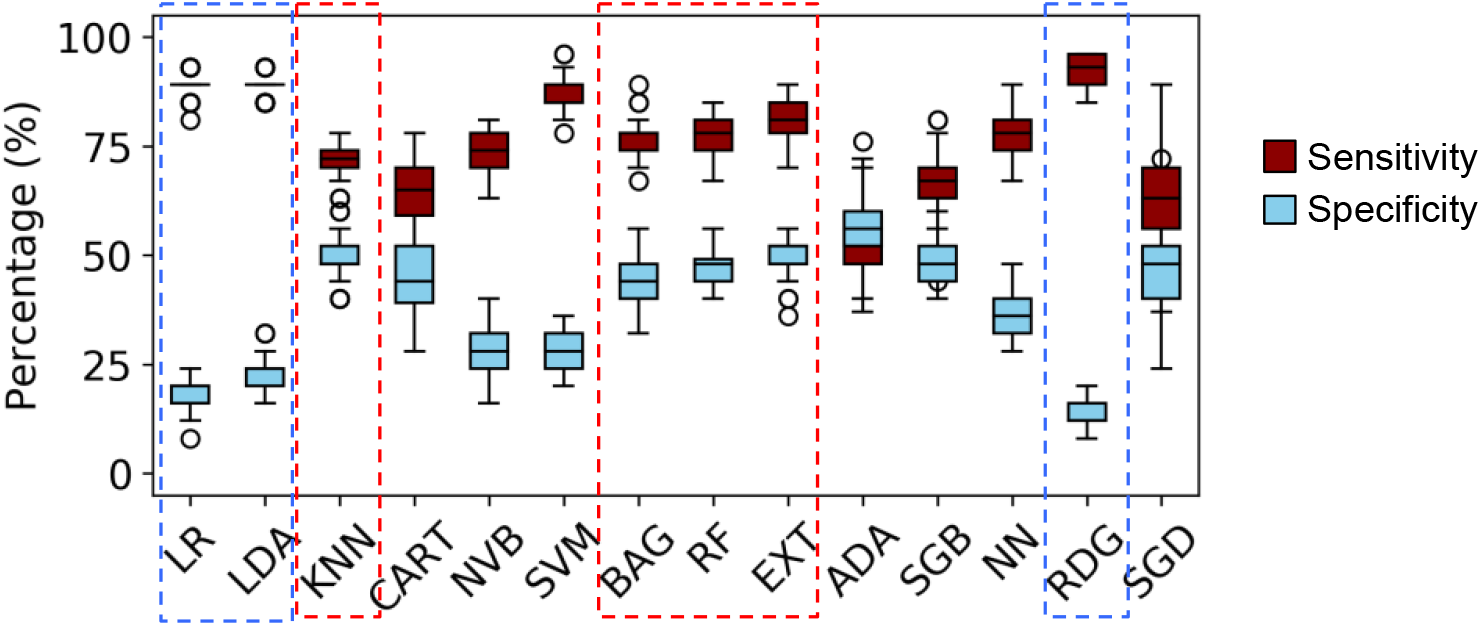
Machine learning model comparison based on repeated stratified 10-fold cross-validation analysis of the LDA-transformed training set. Boxplot of machine learning performance across different models tested shown as sensitivity and specificity based on 100 repeated 10-fold cross-validation. Red dashed boxes indicate the top four models with the highest performance, and blue dashed boxes indicate the top three most sensitive models. **Sensitivity**: proportion of selecting true osteosarcoma; **specificity**: proportion of selecting non-osteosarcoma (Healthy, Other neoplasia, and Non-neoplasia).

The top-performing algorithms were chosen based on their sensitivity and specificity values for further validation (**Fig. 6**). The chosen top four algorithms (KNN, RF, BAG, and EXT) were then retested individually, by the Majority Rule voting approach, and by all-or-none calling method using 10-fold cross-validation with 10 randomized iterations and summarized based on their predictive power, shown as positive (PPV) and negative (NPV) predictive values (**Fig. 6A**). LR, LDA, and RDG were chosen as the three most sensitive algorithms and retested for their predictive power (**Fig. 6B**). We then utilized the post-treatment osteosarcoma samples (n=24; **Table S1**) to evaluate the ability of the “osteosarcoma-detectable” signature to predict the presence of minimal residual disease after treatment, and its relationship to event-free survival outcomes (*i.e.,* duration of remission, **Table S1**). The data were analyzed to form Kaplan-Meier survival curves using the top-four performing algorithms and the three most sensitive algorithms (**Figs. 7A-B**). For this analysis, post-treatment osteosarcoma samples that were classified as “osteosarcoma” by all four models (either best performing based on sensitivity and specificity (**Fig. 5**, red dashed lines) or most sensitive (**Fig. 5**, blue dashed lines)) were considered to be “osteosarcoma – detectable”, and post-treatment osteosarcoma samples that received another classification by one or more of the four models were considered to be “osteosarcoma – NOT detectable”. Event-free survival (time to disease progression) was then compared between dogs whose post-treatment samples were considered “osteosarcoma - detectable” and dogs whose post-treatment samples were considered “osteosarcoma – NOT detectable”. **Fig. 7A** shows that, using the top-four performing models, dogs with post-treatment samples predicted as “osteosarcoma - NOT detectable” had extended event-free survivals (median disease-free interval of 371.1 days) compared to dogs with post-treatment samples predicted as “osteosarcoma-detectable” (median disease-free interval of 149.0 days). The hazard ratio of “NOT detectable” versus “detectable” at 15 months was 2.252, p = 0.1675 (CI: 0.8557 – 5.927). **Fig, 7B** shows that, using the top-three most sensitive models, dogs with post-treatment samples predicted as “osteosarcoma - NOT detectable” had extended event-free survivals (median disease-free interval of 722 days) compared to dogs with post-treatment samples predicted as “osteosarcoma-detectable” (median disease-free interval of 215 days). The hazard ratio at 15 months of “NOT detectable” versus “detectable” was 3.066, p = 0.0398 (CI: 1.054 – 8.922).

**Figure 6.**
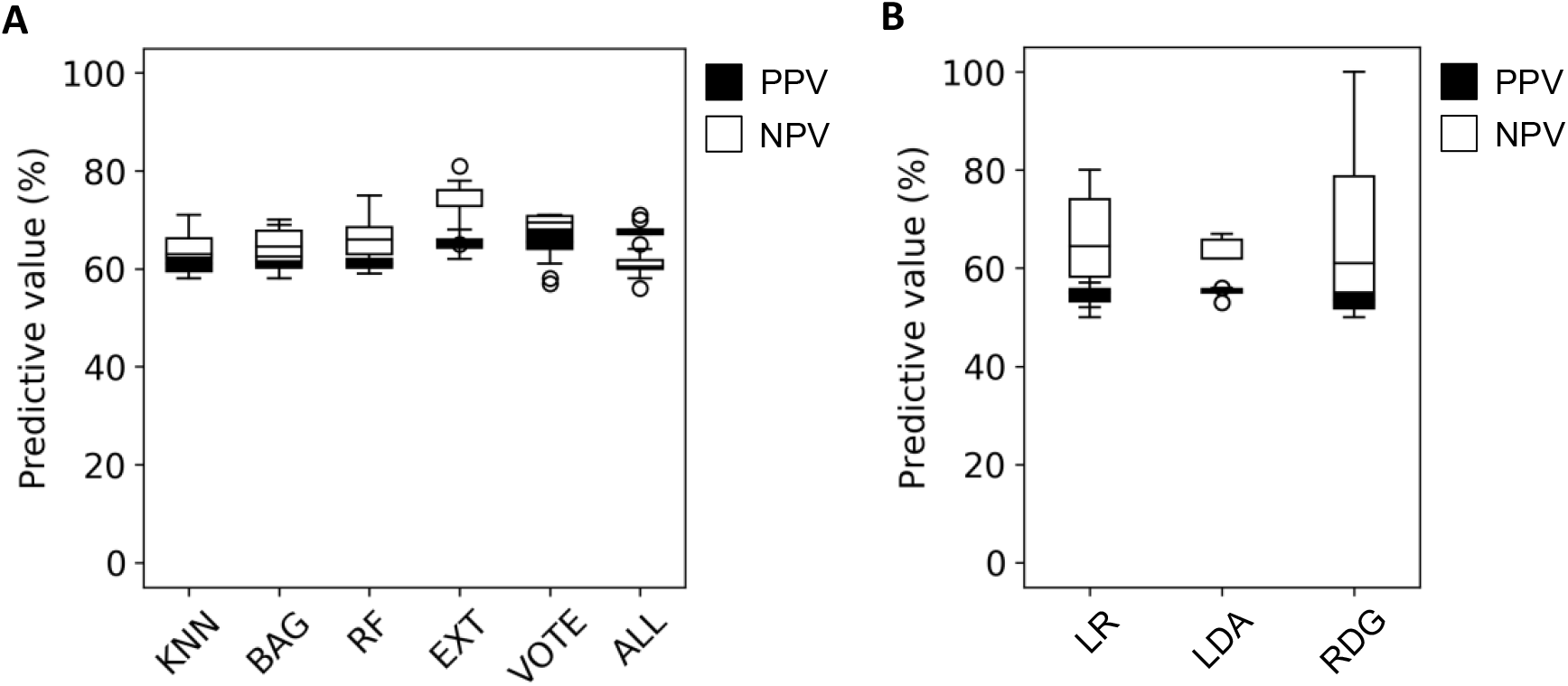
Positive predictive value and negative predictive values. Positive predictive (PPV) and negative predictive (NPV) values (shown as %) based on 10 iterations of shuffled 10-fold cross-validation for (**A**) the top four learning models (KNN, BAG, RF, and EXT) and (**B**) the three most sensitive models (LR, LDA, and RDG). PPV and NPV are also shown for a combined prediction of the top four models shown in A based on the Majority Rule voting (VOTE) and by all-or-none prediction, where Osteosarcoma is called only if agreed by all models (ALL), otherwise the prediction is called as non-Osteosarcoma.

**Figure 7.**
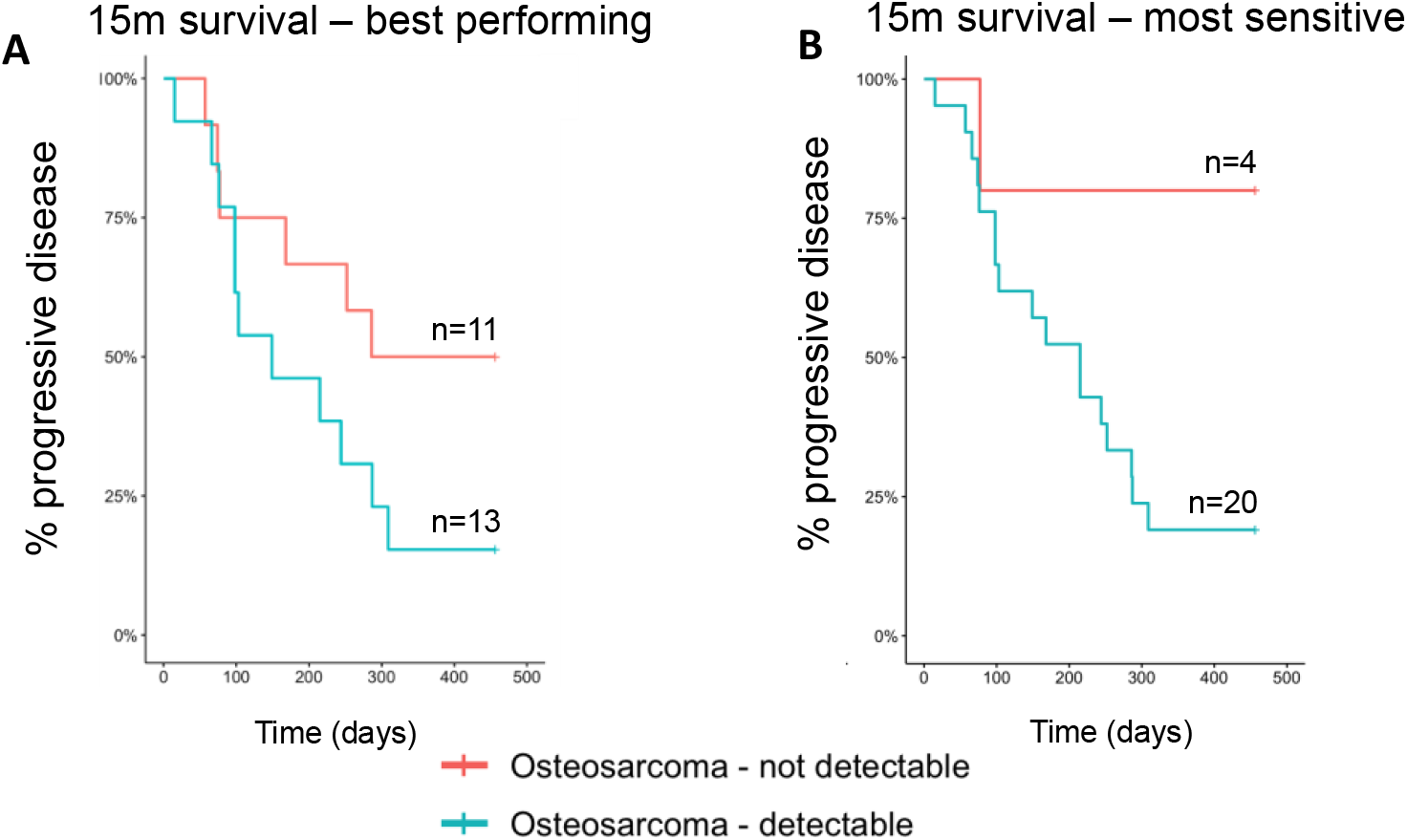
Machine learning models predict presence minimal residual disease in canine osteosarcoma. Post-treatment samples (test set) from dogs with osteosarcoma (n=24) were classified as “osteosarcoma – detectable” or “osteosarcoma – not detectable” based on predictions from (A) the four best performing machine learning models (KNN, BAG, RF, and EXT), or (B) the three most sensitive machine learning models (LR, LDA, RDG). Kaplan-Meier survival curves demonstrating time to relapse for subset of dogs with osteosarcoma with available survival data, comparing those whose post-treatment samples were classified as “osteosarcoma – detectable” with those whose post-treatment samples were classified as “osteosarcoma – not detectable”, (A) p = 0.1001; (B) p = 0.0398.

## Discussion

Non-invasive tests that inform prognosis and longitudinal remission status remain a persistent unmet need for patients with osteosarcoma. In this study, we identified a gene signature associated with prognosis in canine osteosarcoma using a novel xenograft model and bioinformatics pipeline ^17^. We hypothesize that the 5-gene exosomal biomarker signature was associated with prognosis following treatment in dogs with osteosarcoma due to the potential detection of microscopic metastatic disease. The data demonstrate the robustness of our novel xenograft and bioinformatics platform to identify biomarkers for biologically or prognostically significant conditions.

Osteosarcoma, the most common primary tumor of bone, exhibits heterogeneous biological behavior ^3–5^; some tumors are extremely aggressive and unlikely to respond to conventional approaches, whereas others have more variable outcomes and may not require as aggressive treatment protocols. However, stratifying tumors based on aggressiveness is challenging in the clinical setting without suitable biomarkers. Previously identified transcriptional programs that predict tumor behavior and inform prognosis for osteosarcoma patients include the gene cluster expression summary score, or GCESS ^3,39^. The GCESS methodology overcomes the challenge of tumor heterogeneity by identifying coordinately regulated transcripts that provide a cleaner signal, resulting in a biomarker set with greater sensitivity and specificity than that afforded by single biomarkers. However, the GCESS technique requires invasive tissue biopsies ^21^, so it has not been widely adopted in practice; additionally, its utility to monitor minimal residual disease is unknown. Therefore, non-invasive tests that inform prognosis and longitudinal remission status remain a persistent unmet need for patients with osteosarcoma. Development of a prognostic test could allow for patient stratification and development of improved personalized approaches in future clinical research, so that treatment plans minimize risk and maximize benefit, ultimately aiding in the development of new therapies optimized for relative risk.

Extracellular vesicles, specifically exosomes, have great potential to be powerful diagnostic tools. In particular, they are stable in biological fluids, can be efficiently isolated, and contain cargo that is significantly associated with different disease states ^9–11^. The discovery of exosomes and their role in transferring genetic information between cells has sparked interest in utilizing these extracellular vesicles in the discovery of key genes promoting tumor progression ^35,36,40–42^. Indeed, osteosarcoma extracellular vesicle-educated MSCs were shown to promote tumor growth and metastasis in an orthotopic xenograft mouse model of osteosarcoma ^37^. Hoshino et al. utilized exosomal proteins to characterize a variety of tumors, including osteosarcoma; however, the number of included osteosarcoma extracellular vesicle samples was relatively small (7 tumor tissue and 5 plasma extracellular vesicle samples). Given the extensive heterogeneity exhibited by osteosarcoma, thorough characterization using exosomal proteins might be challenging ^43^.

Despite the potential clinical utility of exosomes, identifying cargo originating from diseased cells (the “signal”) from the background of normal exosomes (the “noise”) is still a major obstacle that precludes wide use of exosomes in clinical laboratory medicine. Even in the case of cancer where tumor cells release more exosomes than normal cells, the number of exosomes produced by the tumor is dwarfed by the exosomes produced by the patient’s normal cells, masking all but the strongest tumor-derived exosome signals. Therefore, additional steps, such as sorting by flow cytometry or immunomagnetic enrichment with antibodies or tumor markers, are often undertaken to enrich specifically for cancer-associated exosomes ^44,45^.

The discovery platform described herein, utilizing canine osteosarcoma as an example, allows virtually complete separation of tumor-derived exosomal mRNA cargo and normal cell-derived exosomal mRNA cargo using osteosarcoma xenograft models and an innovative bioinformatics pipeline. Essentially, the mouse model filters the “noise” from the system and helps define tumor and host responses individually. The data from the xenograft models suggest that osteosarcoma-derived exosomes modify the metastatic niche, and host-derived exosomes create a window to understand the host response to the presence of tumors. Utilizing this platform, we developed a 5-gene signature associated with “presence of canine osteosarcoma,” which was further validated in the relevant target species (dogs). The role that these mRNAs play in exosome biosynthesis or in intercellular communication is unclear. It is possible that when exosomes are taken up by cells at distant sites, these mRNAs could be translated in the target cells and contribute to molding the metastatic niche, potentially by immune modulation ^46,47^. On the other hand, they might represent mRNAs that are eliminated from the tumor cells via exosomes because they are toxic when present in high abundance. Nevertheless, we determined that these five mRNAs would provide the foundation for an “osteosarcoma-detectable” signature that would be diagnostically useful. We decided to test the hypothesis that the “osteosarcoma-detectable” signature could be used in a machine learning environment to establish the presence of microscopic, minimal residual disease in dogs with osteosarcoma after surgery +/− chemotherapy. Specifically, machine learning was applied to post-treatment serum samples obtained from dogs with osteosarcoma as a test-set for detection of minimal residual disease following treatment. The “osteosarcoma – not detectable” group had extended event-free survivals compared with the “osteosarcoma – detectable” group using either the top-performing or the most sensitive models, suggesting that the test could detect the presence of osteosarcoma cells (*i.e.*, minimal residual disease) to prognosticate survival after initial treatment. Additional variables with the potential to introduce bias (including random assignment of samples to groups) were compared, and none generated a signature that resulted in robust cross-validation after training.

We hypothesize that dogs where the “osteosarcoma-detectable” signature was present after treatment had a shorter event-free survival due to the presence of minimal residual disease. Additional work is needed to validate the gene signature in an independent set of canine osteosarcoma serum samples. However, the potential to detect minimal residual disease using this signature suggests that it might have utility in the clinical setting for determining prognosis after treatment. This biomarker could be applied after surgery and/or the first round of conventional chemotherapy to guide the subsequent treatment of dogs with osteosarcoma and to alter the course of therapy as needed.

In conclusion, our data support the application of a novel platform consisting of an osteosarcoma xenograft model and bioinformatics pipeline for discovery of prognostically significant, species-specific mRNAs. Moreover, our results document the utility of machine learning algorithms to enhance applicability of these mRNAs to address unmet medical needs, such as sensitive detection of minimal residual disease. Specifically, for this study we identified and validated a 5-gene signature associated with the presence of osteosarcoma in dogs. We further determined that this 5-gene signature obtained from serum exosomes, without the need for more invasive testing, was associated with prognosis, presumably due to the detection of minimal residual disease in dogs with osteosarcoma following treatment. This exosomal 5-gene signature could be applied to clinical veterinary practice and a comparable signature uncovered using our platform could be investigated in human osteosarcoma. Species-appropriate signatures would allow for stratification of dogs and humans with osteosarcoma to minimize risk and maximize benefit of treatment, ultimately aiding in the development of novel therapies.

## Acknowledgments

The authors would like to thank Mitzi Lewellen for assistance with the mouse work, Aaron Sarver and Don Bellgrau for manuscript review and discussion, Mike Farrar, Daniel Vallera, Shruthi Naik, and Steven J. Russell for assistance with funding, and M. Gerard O’Sullivan and Ingrid Cornax for their assistance with mouse pathology. Additionally, we would like to thank Jong-Hyuk Kim, Ashley Schulte, Taylor DePauw, and Erin Dickerson for laboratory support and discussion, Jonah Cullen for assistance with figures, and Dayane Alcantara for technical assistance. Finally, we thank Dr. Holly Borghese of the OSU CVM Veterinary Biospecimen Repository and Blue Buffalo Veterinary Clinical Trials Office (BBVCTO) and Kathleen M. Stuebner, Sara Pracht, Kelly Bergsrud, Andrea Chehadeh, Amber Winter, and Donna Groschen of the Clinical Investigation Center (CIC) of the University of Minnesota for their expertise in sample procurement.

## Ethics Approval and Consent to Participate

This study did not involve the use of human participants, human data, or human tissue (except where commercially sourced); therefore, ethics approval for human participation was not required. All procedures involving animal subjects were approved by the Institutional Animal Care and Use Committees of The University of Minnesota (protocols 0802A27363, 1101A94713, 1312-31131A, 1504-32486A, 1702-34548A, 1803-35759A, and 2003-37952A) and The Ohio State University (protocols 2010A0015-R2 and 2018A00000100).

## Author Contribution Statement

K.M.M., A.J.D., A.K., M.C.S., T.D.O., B.A.B., S.S., and J.F.M. performed study conceptualization and design; K.M.M., A.J.D., A.K., M.C.S., A.R.O., D.C.G., H.T., C.A., K.W., A.M., B.A.B., S.S., and J.F.M. contributed to methodology; K.M.M., A.J.D., A.K., M.C.S., A.R.O., D.C.G., H.T., C.A., K.W., A.M., L.J.M., B.A.B., and J.F.M. performed data curation and investigation; K.M.M., A.J.D., A.K., M.C.S., G.R.C., B.A.B., and J.F.M. performed formal analyses; K.M.M., A.J.D., A.K., M.C.S., A.R.O., D.C.G., G.R.C., B.A.B., S.S., and J.F.M. contributed to data visualization; K.M.M., J.M.F., W.C.K., B.A.B., S.S., and J.F.M. contributed to resources; K.M.M., A.J.D., T.D.O., L.G.S., B.J.W., B.A.B., S.S., and J.F.M. contributed to funding acquisition; A.K. provided software; and B.A.B., S.S., and J.F.M. performed supervision and project administration; K.M.M., A.J.D., A.K., M.C.S., and J.F.M. performed writing of the original manuscript draft. All authors performed review and editing of the manuscript. All authors read and approved the final paper.

## Data Availability Statement

Sequencing data will be deposited in GenBank/GEO and all other data are available from the corresponding author on reasonable request.

