## Supplemental for "Development of an exosomal gene signature to detect residual disease in dogs with osteosarcoma using a novel xenograft platform and machine learning"

**Supplemental material legends**

**Supplemental Table S1.** Machine learning algorithms classified post-treatment osteosarcoma samples as either "osteosarcoma – detected" or "osteosarcoma – not detected". Prediction summaries of post-treatment samples (test set) by "best performing" machine learning models (KNN, BAG, RF, EXT).

**Supplemental Table S2.** Machine learning algorithms classified post-treatment osteosarcoma samples as either "osteosarcoma – detected" or "osteosarcoma – not detected". Prediction summaries of post-treatment samples (test set) by "most sensitive" machine learning models (LR, LDA, RDG).

**Supplemental Figure 1.** Exosome production by osteosarcoma cells is positively correlated with tumor aggressiveness. Cultured dog osteosarcoma cell lines, OSCA-32 and OSCA-40, were stained for tetraspanins CD63, CD9, and CD81 that are enriched in exosomes.

**Supplemental Figure 2.** Physical characterization of serum exosomes. NanoSight particle tracking analysis of triplicate samples serum from healthy control (A) and osteosarcoma-positive (B) representative dogs showing modal diameters of approximately 130 nm.

**Supplemental Figure 3.** Osteosarcoma exosome uptake by fibroblasts and endothelial cells. (A) A CD81-GFP fusion vector was stably transfected into OSCA40 canine osteosarcoma cells. Expression of the fusion protein (green) was determined via confocal immunofluorescent microscopy. Blue staining is the nucleus. (B & C) CD81-GFP tagged exosomes were purified from OSCA40 cells and added to primary cultures of human pulmonary fibroblasts (B) or human pulmonary endothelial cells **(**C**)**. Images of the GFP fluorescence were taken at 0, 1, 2, 4, 8, and 24 hours post treatment with the exosomes. (D & E) Histogram illustrating temporal changes in the number of GFP positive fibroblast and endothelial cells; * $p = 2.62 \times 10^{-6}$; † $p = 2.82 \times 10^{-6}$.

**Supplemental Figure 4.** Machine learning models comparison based on repeated stratified 10-fold cross-validation analysis of the LDA-transformed training set. (A) Classification accuracy of the Healthy group by KNN, BAG, RF, and EXT learning models based on cross-validation analysis. (B) Representative prediction summary of the training set by the top four machine learning models (KNN, BAG, RF, and EXT) based on a 10-fold cross-validation.

**Supplemental Figure 5.** Machine learning performance after data randomization based on repeated 10-fold cross-validations. (A) Original data. (B) Randomized data. (C) Sample distribution was equalized by adding 100 of simulated data points to each group. (D) Randomized equalized data. T-test analysis was performed between original and randomized data. *p = 0.049; ***p < 0.0001

### Supplemental Table S1

| Sample | Diagnosis | Post-treatment classification | Days till relapse |
|---|---|---|---|
| Dog 01 | Osteosarcoma | Osteosarcoma - NOT detectable | 518 |
| Dog 07 | Osteosarcoma | Osteosarcoma - detectable | 244 |
| Dog 11 | Osteosarcoma | Osteosarcoma - NOT detectable | 74 |
| Dog 13 | Osteosarcoma | Osteosarcoma - NOT detectable | 465 |
| Dog 14 | Osteosarcoma | Osteosarcoma - detectable | 215 |
| Dog 15 | Osteosarcoma | Osteosarcoma - NOT detectable | 252 |
| Dog 16 | Osteosarcoma | Osteosarcoma - NOT detectable | 286 |
| Dog 17 | Osteosarcoma | Osteosarcoma - detectable | 98 |
| Dog 20 | Osteosarcoma | Osteosarcoma - NOT detectable | 77 |
| Dog 23 | Osteosarcoma | Osteosarcoma - NOT detectable | 168 |
| Dog 26 | Osteosarcoma | Osteosarcoma - NOT detectable | 57 |
| Dog 29 | Osteosarcoma | Osteosarcoma - NOT detectable | 472 |
| Dog 32 | Osteosarcoma | Osteosarcoma - NOT detectable | 722 |
| Dog 35 | Osteosarcoma | Osteosarcoma - detectable | 98 |
| Dog 36 | Osteosarcoma | Osteosarcoma - detectable | 287 |
| Dog 37 | Osteosarcoma | Osteosarcoma - detectable | 991 |
| Dog 38 | Osteosarcoma | Osteosarcoma - detectable | 309 |
| Dog 39 | Osteosarcoma | Osteosarcoma - detectable | 149 |
| Dog 42 | Osteosarcoma | Osteosarcoma - detectable | 103 |
| Dog 45 | Osteosarcoma | Osteosarcoma - detectable | 76 |
| Dog 48 | Osteosarcoma | Osteosarcoma - NOT detectable | 681 |
| Dog 49 | Osteosarcoma | Osteosarcoma - detectable | 663 |
| Dog 50 | Osteosarcoma | Osteosarcoma - detectable | 15 |
| Dog 53 | Osteosarcoma | Osteosarcoma - detectable | 66 |

**Supplemental Table S1.** Machine learning algorithms classified post-treatment osteosarcoma samples as either “osteosarcoma – detected” or “osteosarcoma – not detected”. Prediction summaries of post-treatment samples (test set) by KNN, BAG, RF, and EXT machine learning models.

### Supplemental Table S2

| Sample | Diagnosis | Post-treatment classification | Days till relapse |
|---|---|---|---|
| Dog 01 | Osteosarcoma | Osteosarcoma - NOT detectable | 518 |
| Dog 07 | Osteosarcoma | Osteosarcoma - detectable | 244 |
| Dog 11 | Osteosarcoma | Osteosarcoma - detectable | 74 |
| Dog 13 | Osteosarcoma | Osteosarcoma - NOT detectable | 465 |
| Dog 14 | Osteosarcoma | Osteosarcoma - detectable | 215 |
| Dog 15 | Osteosarcoma | Osteosarcoma - detectable | 252 |
| Dog 16 | Osteosarcoma | Osteosarcoma - detectable | 286 |
| Dog 17 | Osteosarcoma | Osteosarcoma - detectable | 98 |
| Dog 20 | Osteosarcoma | Osteosarcoma - NOT detectable | 77 |
| Dog 23 | Osteosarcoma | Osteosarcoma - detectable | 168 |
| Dog 26 | Osteosarcoma | Osteosarcoma - detectable | 57 |
| Dog 29 | Osteosarcoma | Osteosarcoma - detectable | 472 |
| Dog 32 | Osteosarcoma | Osteosarcoma - NOT detectable | 722 |
| Dog 35 | Osteosarcoma | Osteosarcoma - detectable | 98 |
| Dog 36 | Osteosarcoma | Osteosarcoma - detectable | 287 |
| Dog 37 | Osteosarcoma | Osteosarcoma - detectable | 991 |
| Dog 38 | Osteosarcoma | Osteosarcoma - detectable | 309 |
| Dog 39 | Osteosarcoma | Osteosarcoma - detectable | 149 |
| Dog 42 | Osteosarcoma | Osteosarcoma - detectable | 103 |
| Dog 45 | Osteosarcoma | Osteosarcoma - detectable | 76 |
| Dog 48 | Osteosarcoma | Osteosarcoma - detectable | 681 |
| Dog 49 | Osteosarcoma | Osteosarcoma - detectable | 663 |
| Dog 50 | Osteosarcoma | Osteosarcoma - detectable | 15 |
| Dog 53 | Osteosarcoma | Osteosarcoma - detectable | 66 |

**Supplemental Table S2.** Machine learning algorithms classified post-treatment osteosarcoma samples as either "osteosarcoma – detected" or "osteosarcoma – not detected". Prediction summaries of post-treatment samples (test set) by "most sensitive" machine learning models (LR, LDA, RDG).

### Supplemental Figure 1

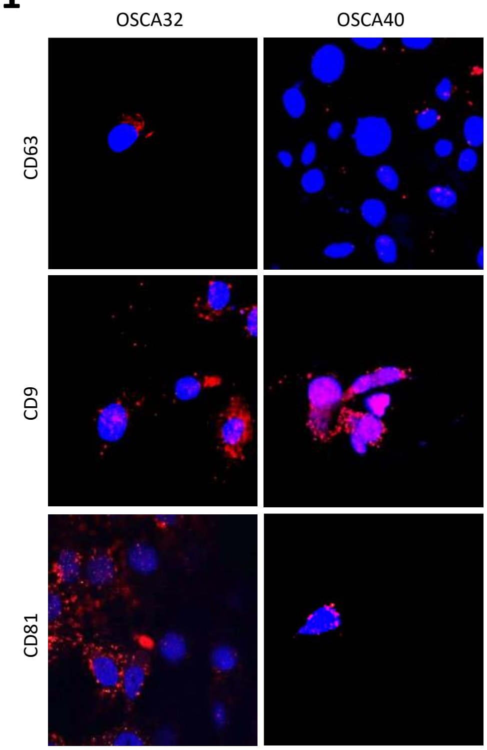


**Supplemental Figure 1.** Exosome production by osteosarcoma cells is positively correlated with tumor aggressiveness. Cultured dog osteosarcoma cell lines, OSCA-32 and OSCA-40, were stained for tetraspanins CD63, CD9, and CD81 that are enriched in exosomes.

### Supplemental Figure 2

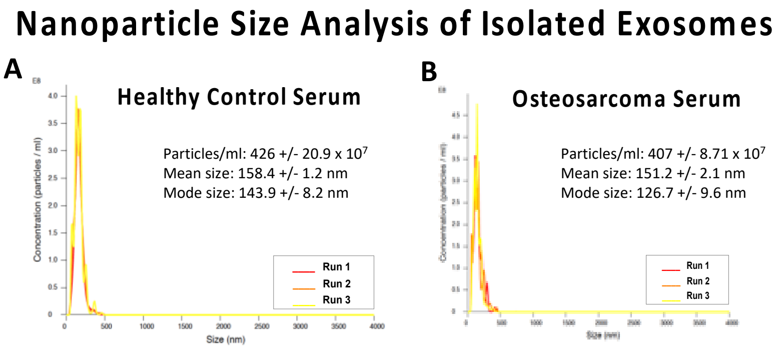


**Supplemental Figure 2.** Physical characterization of serum exosomes. NanoSight particle tracking analysis of triplicate samples serum from healthy control (A) and osteosarcoma-positive (B) representative dogs showing modal diameters of approximately 130 nm.

### Supplemental Figure 3

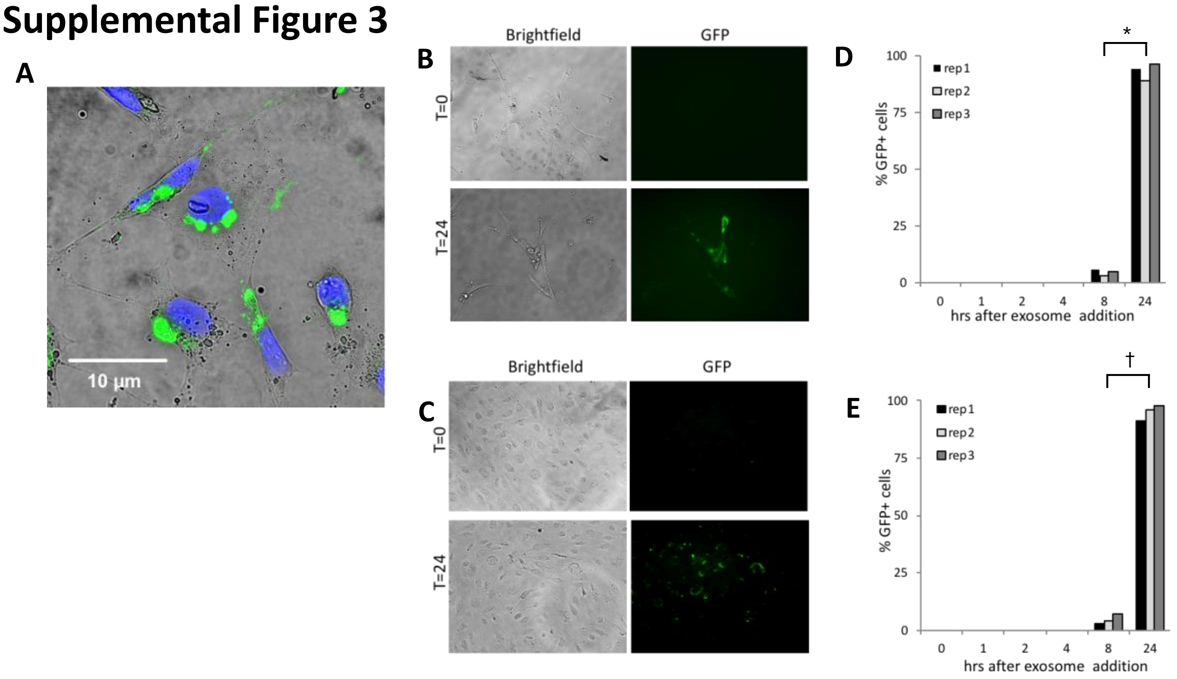


**Supplemental Figure 3.** Osteosarcoma exosome uptake by fibroblasts and endothelial cells. (A) A CD81-GFP fusion vector was stably transfected into OSCA40 canine osteosarcoma cells. Expression of the fusion protein (green) was determined via confocal immunofluorescent microscopy. Blue staining is the nucleus. (B & C) CD81-GFP tagged exosomes were purified from OSCA40 cells and added to primary cultures of human pulmonary fibroblasts (B) or human pulmonary endothelial cells **(**C**)**. Images of the GFP fluorescence were taken at 0, 1, 2, 4, 8, and 24 hours post treatment with the exosomes. (D & E) Histogram illustrating temporal changes in the number of GFP positive fibroblast and endothelial cells; * $p = 2.62 \times 10^{-6}$; † $p = 2.82 \times 10^{-6}$.

### Supplemental Figure 4

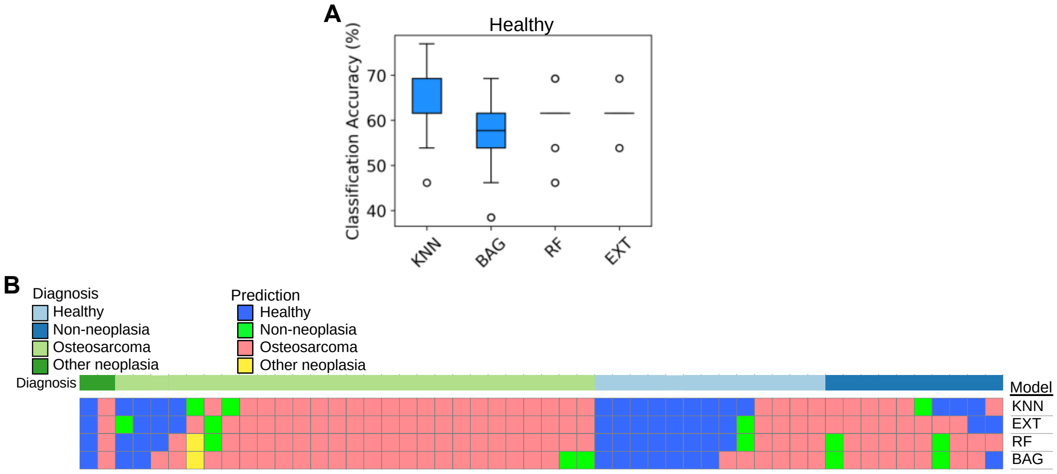


**Supplemental Figure 4.** Machine learning models comparison based on repeated stratified 10-fold cross-validation analysis of the LDA-transformed training set. (A) Classification accuracy of the Healthy group by KNN, BAG, RF, and EXT learning models based on cross-validation analysis. (B) Representative prediction summary of the training set by the top four machine learning models (KNN, BAG, RF, and EXT) based on a 10-fold cross-validation.

### Supplemental Figure 5

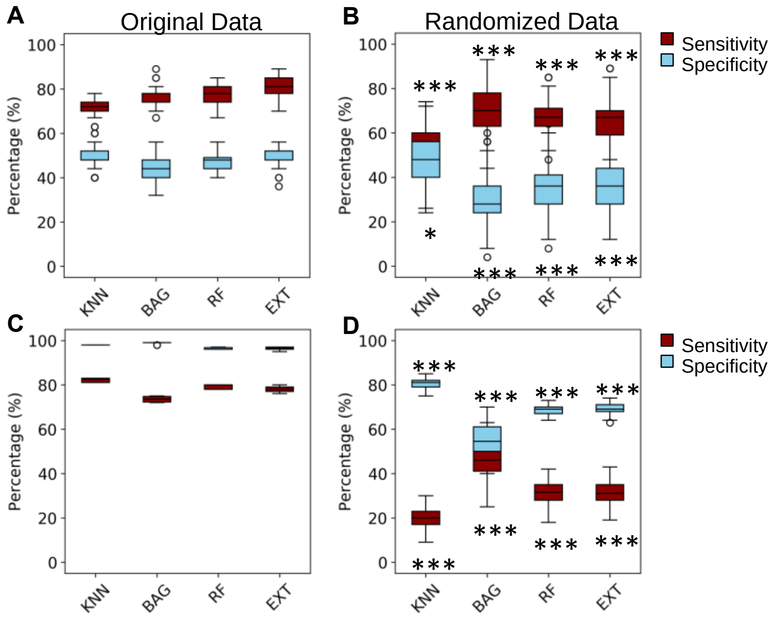


**Supplemental Figure 5.** Machine learning performance after data randomization based on repeated 10-fold cross-validations. (A) Original data. (B) Randomized data. (C) Sample distribution was equalized by adding 100 of simulated data points to each group. (D) Randomized equalized data. T-test analysis was performed between original and randomized data. *p = 0.049; ***p < 0.0001
